# Network Dynamics Theory of Human Intelligence

**DOI:** 10.1101/081695

**Authors:** Aki Nikolaidis, Aron K. Barbey

## Abstract

Scientific discovery and insight into the biological foundations of human intelligence have advanced considerably with progress in neuroimaging. Neuroimaging methods allow for not only an exploration of what biological characteristics underlie intelligence and creativity, but also a detailed assessment of how these biological characteristics emerge through child and adolescent development. In the past 10 years, functional connectivity, a metric of coherence in activation across brain regions, has been used extensively to probe cognitive function; however more recently neuroscientists have begun to investigate the dynamics of these functional connectivity patterns, revealing important insight into these networks as a result. In the present article, we expand current theories on the neural basis of human intelligence by developing a framework that integrates both how short-term dynamic fluctuations in brain networks and long-term development of brain networks over time contribute to intelligence and creativity. Applying this framework, we propose testable hypotheses regarding the neural and developmental correlates of intelligence. We review important topics in both network neuroscience and developmental neuroscience, and we consolidate these insights into a Network Dynamics Theory of human intelligence.

## Introduction

For centuries the nature of human intelligence has motivated considerable research and debate. What mental abilities underlie intelligent behavior and how do they contribute to the expression of genius and creativity? How are these abilities shaped by the environment, cultivated through experience, and represented within the architecture of the human brain? While the precision of scientific theories and methods for investigating these questions has evolved over the years, in recent decades advances in cognitive neuroscience have afforded unprecedented insight into the nature and mechanisms of intelligence and creativity in the human brain. Indeed, the advent of neuroimaging methods has provided an opportunity to study the structural and functional organization of the brain as a window into the architecture of these higher order cognitive capacities in the human mind. Contemporary neurobiological theories have applied neuroimaging methods to establish that individual differences in general intelligence can be localized to a specific network – the fronto-parietal network—whose functions are largely believed to reflect intrinsic and stable computational properties that enable core facets of both intelligence and creativity (Beaty et al., 2014; Barbey, Colom, & Grafman, 2013a; Barbey et al., 2012; Duncan, 2010; Jung & Haier, 2007). Recent evidence from network and developmental neuroscience further demonstrates that static networks undergo both short-term dynamic fluctuations (Beaty, Benedek, Kaufman, & Silvia, 2015; Byrge, Sporns, & Smith, 2014; Deco, Jirsa, & McIntosh, 2011), and long-term changes over the developmental trajectory of the child and adolescent brain (DiMartino et al., 2014; Hutchison & Morton, 2015), and therefore motivate new perspectives about the dynamic (rather than static) and system-wide (rather than singular) network properties that underlie human intelligence and creativity.

In this article, we introduce a cognitive neuroscience framework for understanding the nature and mechanisms of human intelligence, the Network Dynamics Theory, and review evidence to elucidate how functional brain networks and their dynamic properties underlie intelligence and its emergence over childhood and adolescence. According to this framework, intelligence emerges through the process of actively selecting and creating information that in turn modifies the brain’s internal structure and dynamics (Figure 1). The development of these internal dynamics over childhood and adolescence contribute to the maturation of the higher cognitive abilities associated with intelligence (Byrge et al., 2014) and likely creativity. We begin by surveying recent theoretical and experimental advances in network neuroscience that elucidate the static and dynamic brain networks underlying human intelligence and creativity, followed by a review of the neurodevelopmental trajectory of intelligence, and how the maturation of these networks contributes to the formation of both intelligence and creativity in adulthood.

**Figure 1.**
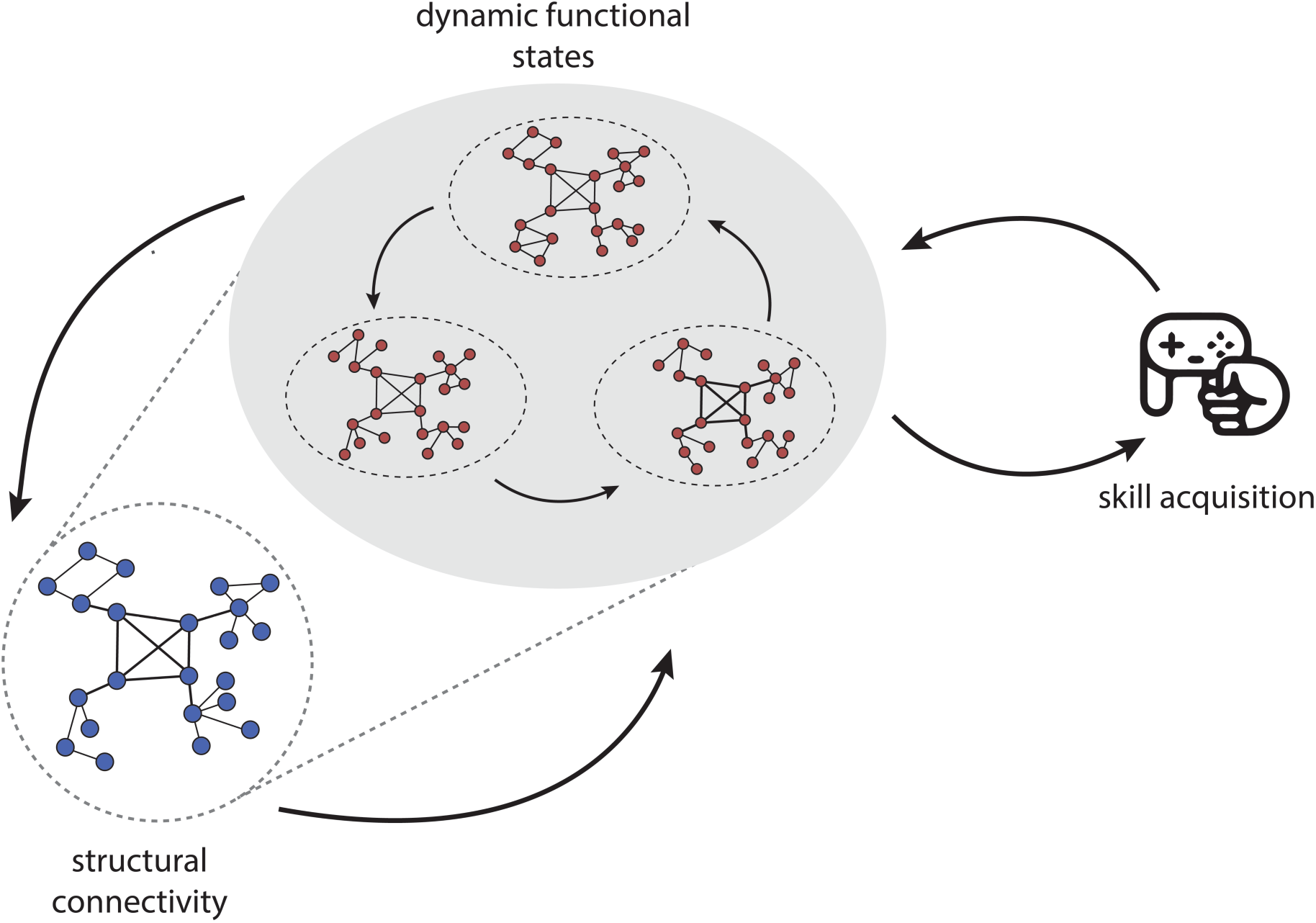
This figure represents how intrinsic and extrinsic forces drive the concurrent development of brain networks and cognitive function (Byrge et al., 2014). Structural brain networks (blue) play a constraining role on the intrinsic brain dynamics of functional networks (red), which in turn modulate the structural networks. We see how this interaction between structural and functional dynamics leads to differences in skill acquisition and response variation. The structure-function relationship constrains the generation of output, motor output in this case, though this could be generalized to cognitive skill performance as well. Sensory inputs deliver error feedback to the brain and new dynamics emerge to generate novel forms of activity. Novel forms of activity are generated and tested for relevant outcomes. Over the course of development this interactive process guides the maturation of structural networks and functional brain dynamics.

### Information Processing Assumptions for Intelligence in the Human Brain

Network Dynamics Theory rests on six principles for information processing in the human brain (Just & Varma, 2007; Newman & Just, 2005). Each of the six principles is well established in the neuroscience literature and motivates predictions about the nature and origins of individual differences in general intelligence.

1. Intelligence is the product of the concurrent activity of multiple brain areas that collaborate in a large-scale cortical network (Barbey et al., 2012; Gläscher et al., 2010). Variation in the degree of synchronization or efficiency of the communication between regions may therefore predict individual differences in task performance (Nikolaidis, Goatz, Smaragdis, & Kramer, 2015).
2. Each cortical area can perform multiple cognitive functions, and conversely, many cognitive functions are performed by more than one area. The diverse functional role of brain regions is evident in literature indicating that regions of the prefrontal cortex serve multiple cognitive functions (Barbey, Colom, & Grafman, 2013b; Duncan, 2010; Miller & Cohen, 2001), and even motor regions play an important role in higher cognition (Nikolaidis et al., 2016; Nikolaidis, Voss, Lee, Vo, & Kramer, 2014; Sabaté, González, & Rodríguez, 2004; Vakhtin, Ryman, Flores, & Jung, 2014). Conversely, the capacity for functional localization across multiple brain regions is well established by the neuroscience literature on cortical plasticity (for a review, see Pascual-Leone, Amedi, Fregni, & Merabet, 2005).
3. Each cortical area has a limited capacity of computational resources, constraining its activity. Evidence indicates, for example, that activity within working memory networks increases with performance gains on the N-Back task and plateaus or decreases as the participant reaches the ceiling of performance (Callicott et al., 1999; Jaeggi et al., 2007). The limited capacity principle has direct implications for individual differences in intelligence. First, it suggests that the amount of resources available or the resource capacity within the neural system varies across individuals, which is supported by evidence demonstrating that individual differences in brain metabolism contribute to cognitive performance (Jung et al., 1999, 2005; Nikolaidis et al., 2016; Paul et al., 2016; Ross & Sachdev, 2004). Secondly, the amount of resources required to perform a task likely differs across individuals due to variations in efficiency (Jaeggi et al., 2007).
4. The topology of a large-scale cortical network changes dynamically during cognition, adapting itself to the functional demands of the task and resource limitations of different cortical areas (Byrge et al., 2014). This principle is supported, for example, by evidence demonstrating that cognitive control networks shift their connectivity in a task dependent manner to dynamically reconfigure brain networks for goal-directed behavior (Cole et al., 2013; Miller & Cohen, 2001). These dynamic network features may therefore contribute to individual differences in goal-directed, intelligent and creative behavior.
5. The communications infrastructure that supports the transfer of information across multiple brain regions is also subject to resource constraints (i.e., bandwidth limitations). This principle is supported by a large body of neuroscience evidence demonstrating that white matter fiber tracts enable the integration of information across broadly distributed cortical networks and that the fidelity of these pathways is critical to general intelligence (Penke et al., 2012). Variation in the degree or quality of the anatomical connections between processing regions may therefore contribute to individual differences in task performance.
6. Neuroimaging measures of cortical activity (e.g., fMRI) provide an index of cognitive workload and computational demand. Extensive neuroscience evidence supports this principle, demonstrating that the amount of cortical activation within a given region increases with computational demands, for example, in sentence comprehension (Röder, Stock, Neville, Bien, & Rösler, 2002), working memory (Braver, Cohen, Nystrom, & Jonides, 1997), and mental rotation tasks (Just, Carpenter, Maguire, Diwadkar, & McMains, 2001).

## Network Dynamics Theory of Human Intelligence

The reviewed operating principles provide the foundation for an interactive system of intrinsic connectivity networks, which together comprise the information processing architecture of human intelligence. Analysis of patterns of functional brain connectivity have revealed statistical dependencies in neural activity across regions, comprising core intrinsic connectivity networks of the brain (see Figure 2; Power, Cohen, et al., 2011), and indicates that these patterns of brain connectivity can be used to predict performance for both high- (Finn et al., 2015) and low-level cognitive processes (Nikolaidis et al., 2015). Functional brain networks are known to fluctuate and evolve over short timescales and are constrained by structural connectivity, which modulate over longer timescales (Byrge et al., 2014; Deco et al., 2011). Critically, functional networks do not relay neural signals, but instead reflect neural communication within an underlying structural network (van den Heuvel & Sporns, 2013). In this way functional networks are an important lens for understanding both low-level processing and a high-level holistic investigation of large-scale networks. Intrinsic connectivity networks are thus characterized by their micro- and macro-level topology. Micro-level topological properties describe local features of the network (e.g., the degree of a target node; Table 1). Macro-level topological properties reflect the large-scale architecture and global organization of the network (e.g., global efficiency; Table 1).

**Table 1.**
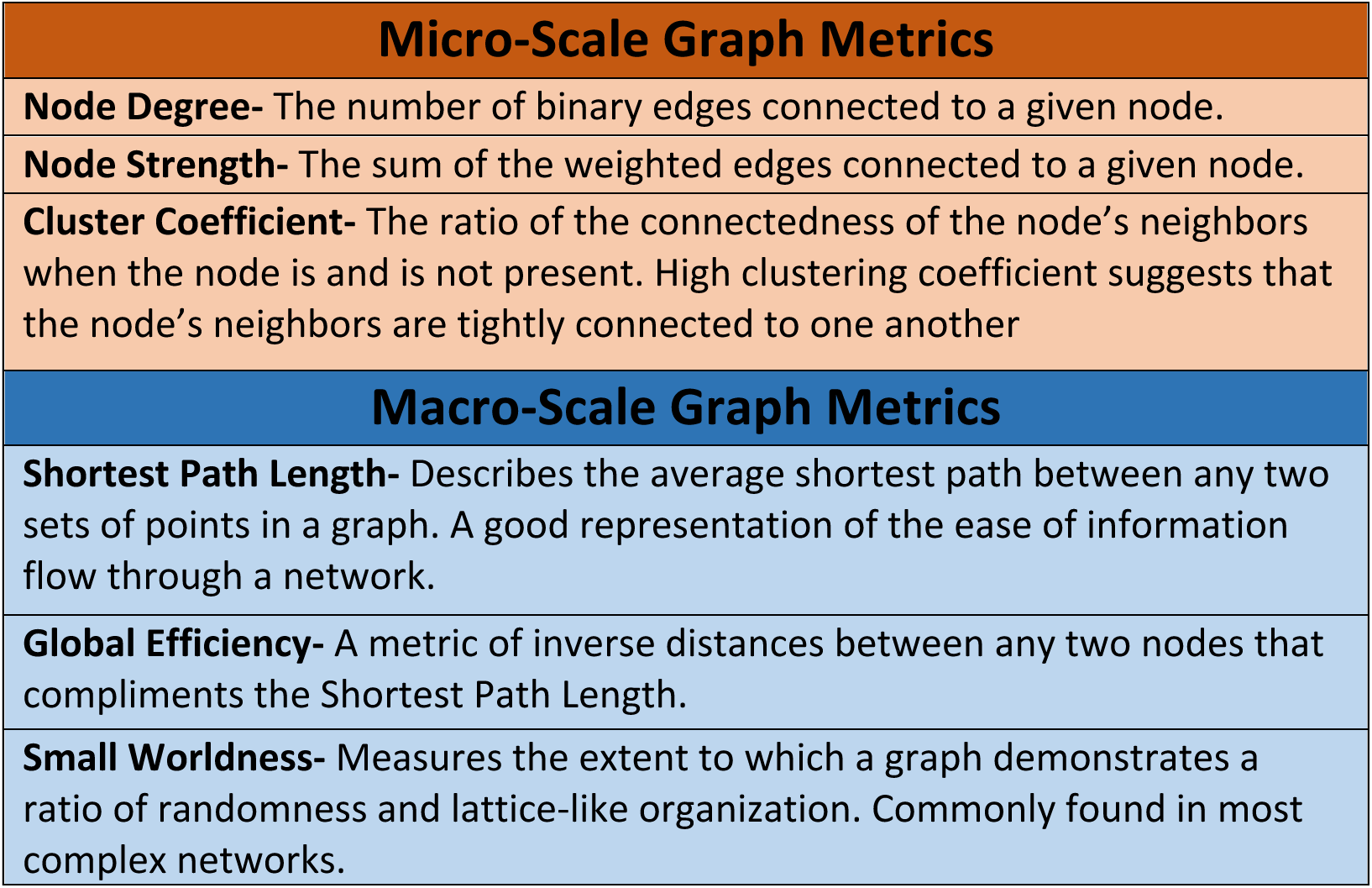
This table summarizes some of the most important micro and macro level graph theoretical measurements of functional network construction (Bullmore & Bassett, 2011; Bullmore & Sporns, 2009; Rubinov & Sporns, 2009). The micro measurements characterize the role of a given node in the network as a whole, while the macro measurements describe various aspects of the construction of the whole network, such as the speed of information transfer.

**Figure 2.**
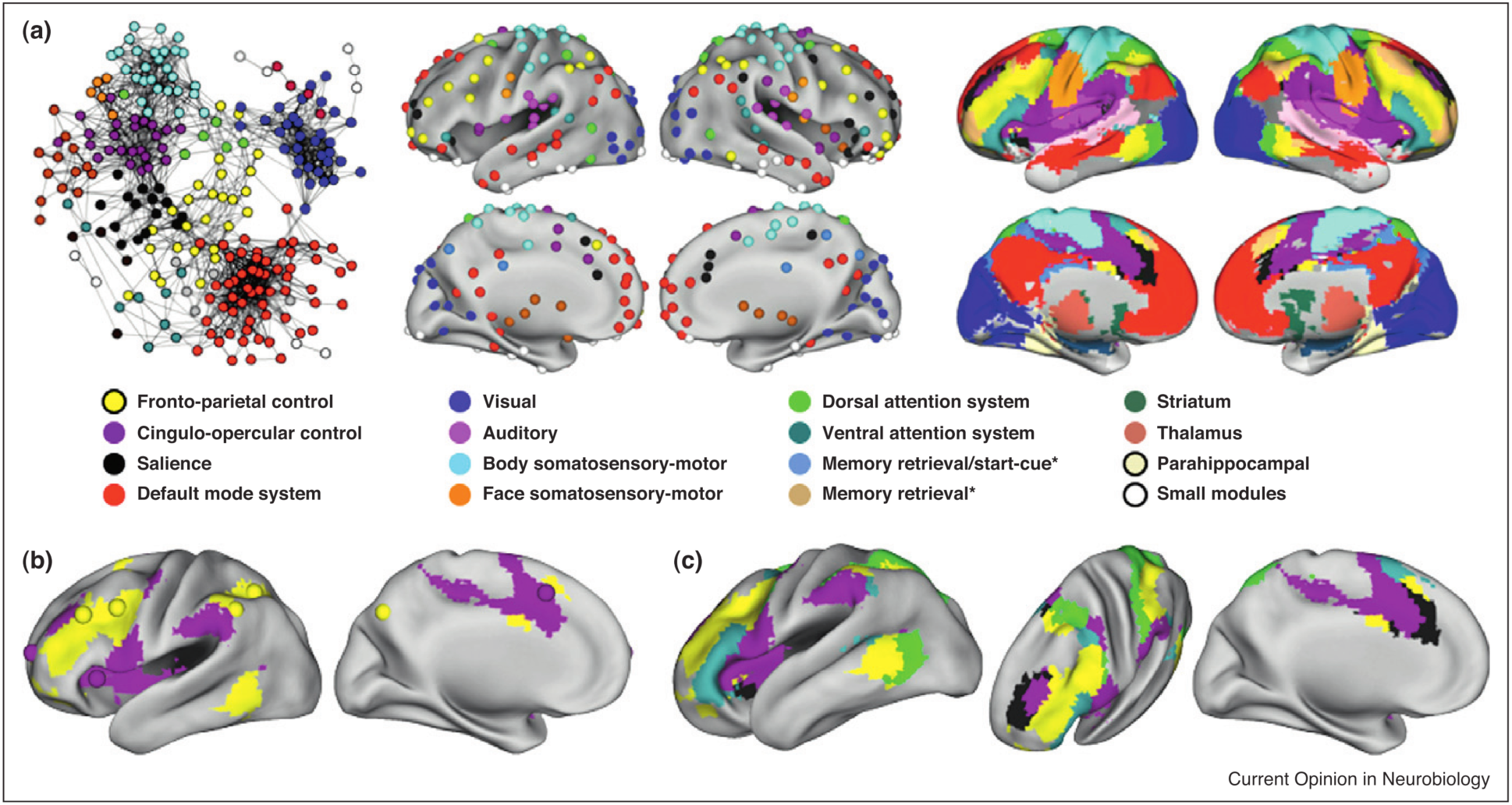
This figure summarizes recent work extracting reliable functional networks based on a large scale meta-analysis of peaks of brain activity for a wide range of motor, perceptual, and cognitive tasks (With permission from Dosenbach et al., 2006; Power & Petersen, 2013). A) The upper left figure represents a graph theoretic embedding of the nodes. Similarity between nodes is represented by spatial distance, and nodes are assigned to their corresponding network by color. The next two sections present the nodal and voxel-wise network distribution in both hemispheres. The bottom panel (B & C) displays a voxel-wise distribution of the cognitive control networks: the frontal parietal network (yellow), the cingulo-opercular network (purple), the dorsal attention network (green), the salience network (black), and the ventral attention network (teal).

Network Dynamics Theory proposes that intelligence fundamentally depends on the learnability of macro-level network structures and dynamics (topological patterns) that emerge from external input. According to this account, intelligent, goal-directed behavior reflects the learner’s capacity to utilize macro-level topological network patterns to process incoming information. This prediction motivates a more precise characterization of the large-scale cortical network properties that underlie human intelligence (from Information Processing Assumption 1). Specifically, hubs are known to play a central role in the formation of macro-level network structures and mediate many of the long-distance connections between brain modules (Figure 3; van den Heuvel & Sporns, 2013). Hub regions, such as the bilateral precuneus, anterior and posterior cingulate cortex, insular cortex, and superior frontal cortex are also known to form a strongly interconnected network of regions (i.e., the rich club network; Figure 3; van den Heuvel, Kahn, Goñi, & Sporns, 2012). Given the range of network and functional roles of these hubs (Information Processing Assumption 2), their associated high computational cost (Information Processing Assumption 3), and high degree of interaction (van den Heuvel et al., 2012), Network Dynamics Theory proposes that the macro-level topological properties of this rich club network play a central role in human intelligence. Specifically, such properties are functionally valuable for integrative information processing and adaptive behavior. For example, network hubs in the fronto-parietal network demonstrate a significant degree of task-specific interactions with a wide variety of cognitive and sensory networks, modulating their connectivity and supporting a diversity of cognitive tasks (Cole et al., 2013). Furthermore, interactions between rich club regions play an important role in determining global efficiency of communication in a network, as demonstrated by evidence indicating that almost 70% of the shortest paths through a whole brain network pass through the rich club (van den Heuvel et al., 2012). Given the role of efficiency of network communication in cognition (Bullmore & Sporns, 2009; Langeslag et al., 2012; Moussa et al., 2011), we propose that the construction of the rich club plays a primary role in the link between functional brain networks and intelligence.

**Figure 3.**
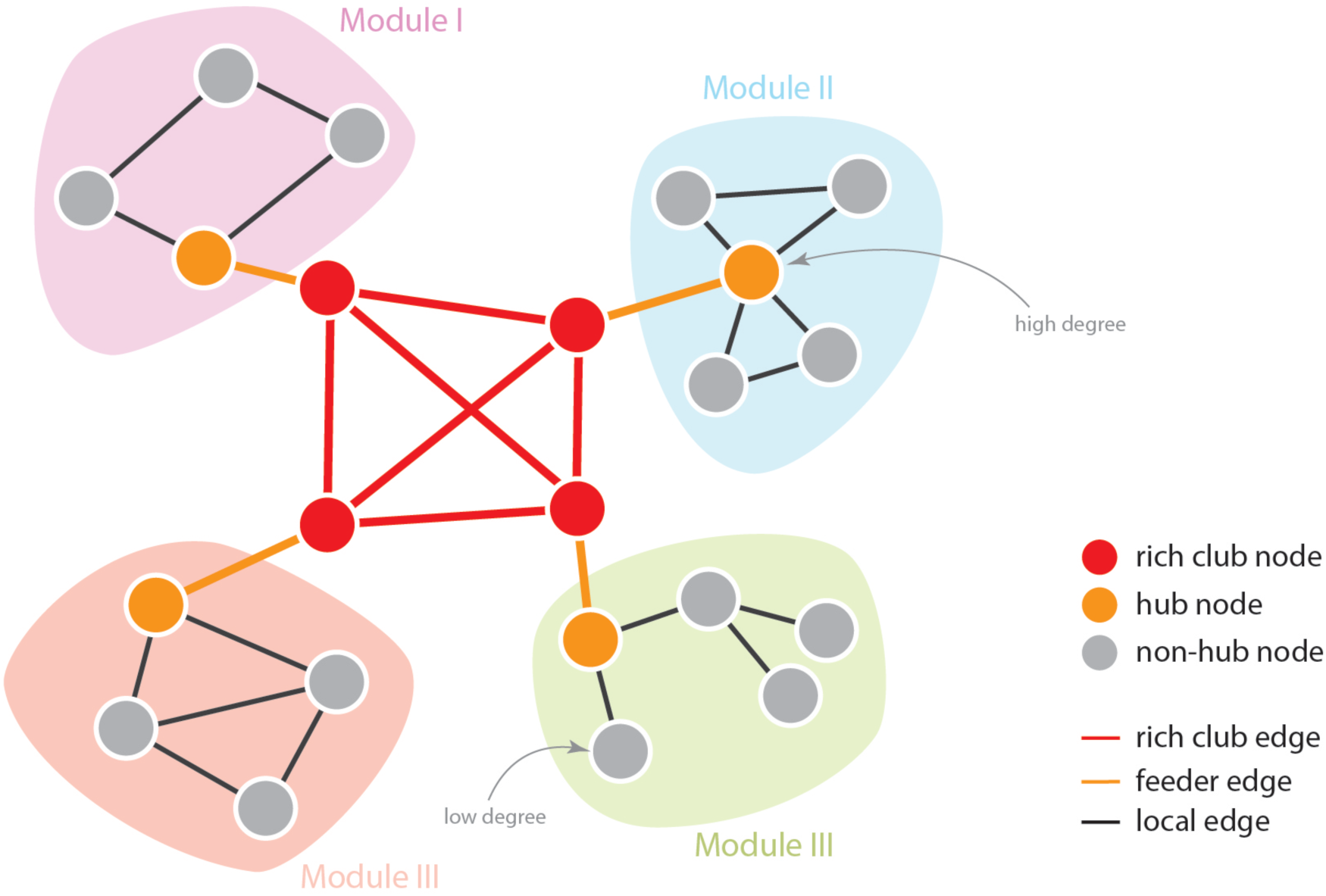
This figure displays a visual summary of basic network structure (van den Heuvel & Sporns, 2013). Each circle is a node and all the connections between them are labeled edges. Nodes of high or low degree, represented as black and gray circles, are those with edges connecting to many or few other nodes respectively. Modules are clusters of nodes with relatively high within-cluster connectivity and low between-cluster connectivity. Among all nodes in the two graphs, the red nodes are designated as hubs given their high-degree and graph centrality, (Harriger, van den Heuvel, & Sporns, 2012). This figure abstractly displays how normal and rich club nodes interact, demonstrating that the rich club nodes not only have high degree, but they also serve as critical way points that enable efficient graph traversal between distant nodes.

As a systematically constrained information-processing organ that has evolved over millennia to maximize computational capacity towards reproductive success, the brain is driven to perform complex computations with a limited budget of resources (Gazzaniga, 2000; Güntürkün, 2005; Roth & Dicke, 2005). Nevertheless, network hubs are sinks for the brain’s metabolic and communication resources. For example, the spatial distance of edges connecting hubs to the rest of the network (i.e., the wiring cost) is greater than the distance of edges connecting more peripheral nodes (i.e., hubs have a high wiring cost) (van den Heuvel et al., 2012). Rich club connections are comparatively costly and their connectivity alone accounts for 40% of the total whole brain communication cost (van den Heuvel et al., 2012). As indicated by Information Processing Assumption 2, each brain region may perform multiple cognitive functions, and this assists with the computational and resource load on the brain. Network hubs are regions that are tightly integrated into single or multiple networks. Furthermore, hubs are known to have higher rates of cerebral blood flow, aerobic glycolysis, and oxidative glucose metabolism (Information Processing Assumption 3). This combination of higher metabolic rate and longer connection distance makes hubs biologically very costly (Crossley et al., 2014). Thus, the high value and high biological cost of hub regions makes them particularly sensitive predictors of individual differences in human intelligence. For example, recent work investigating the role of brain metabolism in cognition has demonstrated that the concentration of NAA, a marker of oxidative metabolism, is a strong predictor of intelligence (Nikolaidis et al., 2016; Paul et al., 2016). Given the brain’s resource constraints, and the cost of using and maintaining these high wiring cost regions, Network Dynamics Theory proposes that the connectivity and activity of these regions may play a particularly important role in cognitive development and the emergence of intelligence. Understanding how these rich club nodes contribute to the development of executive functions is therefore essential to characterizing how the network architecture of the brain shapes intelligence.

### Cognitive Control Functions are Central to Human Intelligence

Cognitive control is a hallmark of human intelligence. The capacity to adaptively re-organize one’s thoughts and actions in accordance with internal goals is an important marker for the development of intelligence. According to Network Dynamics Theory, human intelligence reflects a self-organizing system that adaptively engages multiple brain networks to support goal-directed, purposeful behavior. To successfully perform a particular task, mental operations must be selected to achieve that specific task out of an infinite number of possible tasks and corresponding mental operations (Duncan, 2010). The process of selecting and implementing behavior-guiding principles that enable goal achievement is the central question of cognitive control. At least three signals may be defined that cognitive control regions should display across a wide variety of tasks (Figure 4; Power & Petersen, 2013). First, when a subject is given a cue to begin a particular task, control regions must send configuring signals to processors to establish the correct processing strategy needed for the task (the task set). A control region may therefore display start-cue activity as the task set is selected and instantiated. Second, for as long as a subject continues to perform the task, the task set must be maintained. A control region may therefore display sustained activity during task performance. Third, because successful control needs to recognize errors in performance and adjust task set accordingly, a control region may display error-specific activity (Figure 4).

**Figure 4.**
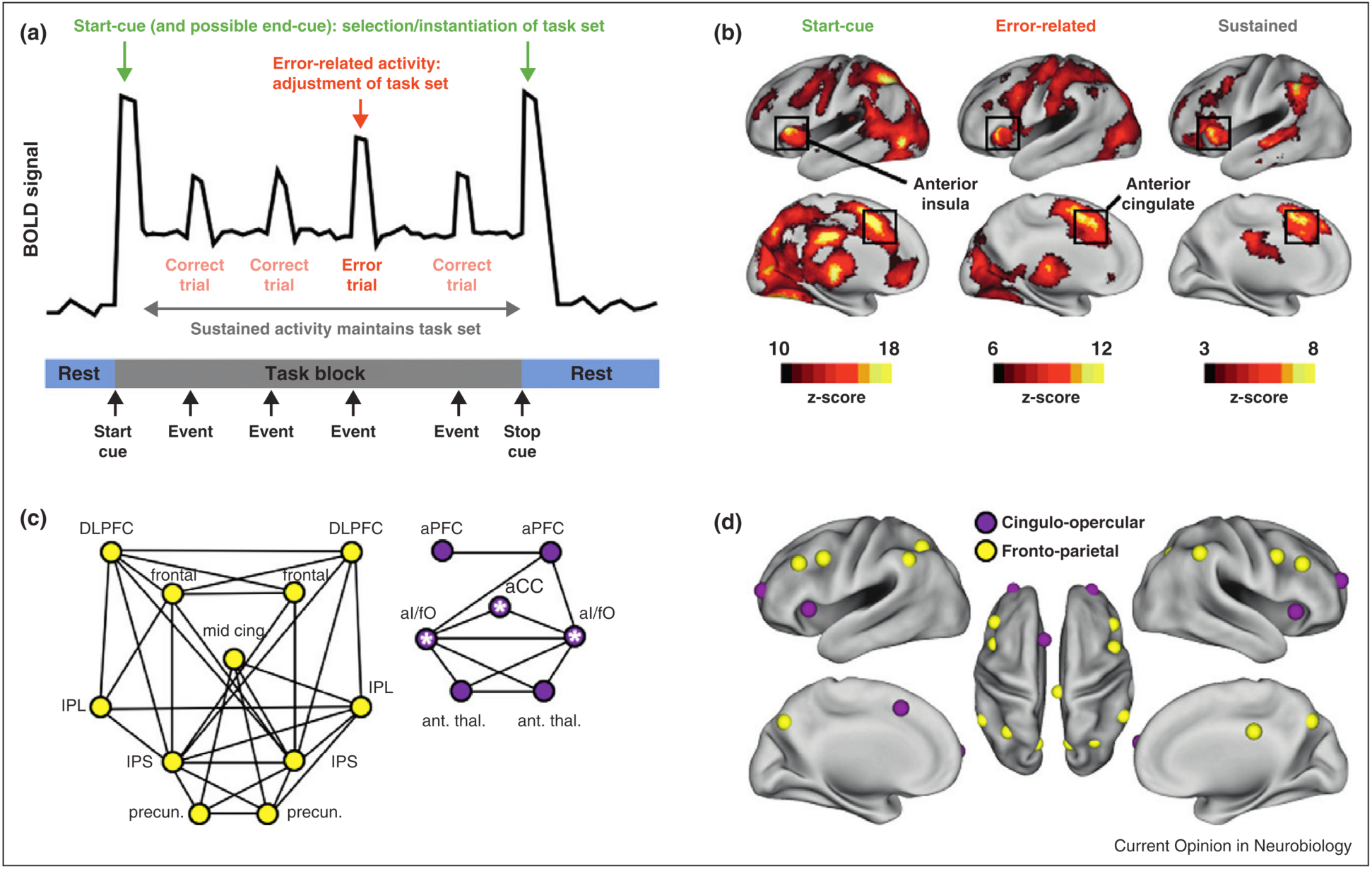
This image represents the brain activity and network contributions to the three cognitive components of cognitive control: Start cue, error related, and sustained activity (Power & Petersen, 2013). A) A hypothetical hemodynamic response function time course from a region that elucidates the response for the start-cue, error-related, and sustained attention components of cognitive control. B) fMRI activity maps that display the distribution of activity sensitive to the start-cue, error signal, and sustained attention. C) A graph that summarizes the connectivity of the frontal parietal network (FPN, yellow) and the cingulo-opercular network (CON, purple). FPN: DLPFC-dosal lateral prefrontal cortex, IPL- inferior parietal lobe, IPS- Intra-parietal sulcus. CON: aPFC- anterior prefrontal cortex, al/fO- anterior insula/frontal operculum, aCC- anterior cingulate cortex. D) The anatomical mappings of the two cognitive control networks abstractly represented in C.

Specific intrinsic connectivity networks are known to support cognitive control processes, including the fronto-parietal network and the cingulo-opercular network (Dosenbach, Fair, Cohen, Schlaggar, & Petersen, 2008). The fronto-parietal network supports moment-to-moment task adjustments and is engaged during start-cue and error-related activity (but does not demonstrate sustained activity during task set maintenance) (Dosenbach et al., 2008). The cingulo-opercular network operates over longer time-scales and is recruited during start-cue, error-related, and sustained activity (Dosenbach et al., 2008). Recent evidence further suggests that cognitive control capacity may be supported by whole-brain network properties. Studies have shown that higher global efficiency of functional brain networks is positively correlated with better cognitive performance (Giessing, Thiel, Alexander-Bloch, Patel, & Bullmore, 2013; van den Heuvel, Stam, Kahn, & Hulshoff Pol, 2009), and that cognitively more demanding tasks may necessitate long-range integrative connections (Kitzbichler, Henson, Smith, Nathan, & Bullmore, 2011). These data suggest that intelligence depends on an integrative network topology (Dehaene & Changeux, 2011). In particular, the rich club of highly-interconnected hub nodes, many of which are in the fronto-parietal network (Cole et al., 2013), are known to support performance in a variety of tasks, especially cognitive control tasks demanding goal-directed thought and behavior. This highlights the value of hubs for the dynamic integrative processes and adaptive behavior that are essential to cognitive control (Crossley et al., 2013, 2014). More recently, Cole and colleagues (2013) found that the fronto-parietal network demonstrates especially high global connectivity across a wide variety of tasks. This finding suggests that global connectivity of specific control regions may be important for cognitive control capacity and would allow for a mechanism by which specific control regions can access and influence other relevant networks (such as sensory-motor networks involved in task-relevant processing) to adaptively monitor and regulate ongoing behavior (Dehaene, Kerszberg, & Changeux, 1998; Miller & Cohen, 2001).

Dynamic Variability of Functional Brain Networks in Human Intelligence

While the topology of the brain’s structural connectivity plays an important role in constraining brain connectivity (Figure 1; Information Processing Assumption 5), the interactions between regions demonstrate significant variability over shorter time scales (Information Processing Assumption 4) (Deco et al., 2011). An emerging area of research in network neuroscience investigates how interactions between cortical areas enable human intelligence (Cole et al., 2013; Hampshire, Highfield, Parkin, & Owen, 2012). This research indicates that the interaction among brain regions is dynamic – the system adaptively configures and reconfigures itself in light of changes in processing demands and inherent limitations in available computational resources. While regions with highly stable pairwise connectivity may demonstrate such strong connectivity as the result of direct callosal fiber connections (e.g., in the case in bilateral homologies), many higher-order regions demonstrate greater variability in functional connectivity and tend to be involved in a greater range of functions (Deco et al., 2011).

On the basis of these findings, Network Dynamics Theory proposes that dynamic variability in functional connectivity is critical for the diverse range of processing involved in intelligence, and recent work on dynamic brain connectivity has shed light onto how these dynamic networks relate to static functional connectivity networks and cognition. Variability in functional interactions between nodes gives rise to a large set of functional network states that are strongly fluctuating over time, and which may differ from commonly defined static networks (Hutchison et al., 2013). In fact, well-defined static networks, such as the default mode network (DMN), actually pass through multiple meta-stable states (Allen et al., 2014), and these short-lived functional connectivity states are reproducible across subjects (Allen et al., 2014; Liu, Chang, & Duyn, 2013). The fronto-parietal network is made flexible through its composition of hubs that rapidly modulate their pattern of global functional connectivity according to task demands (Cole et al., 2013). This work also demonstrates the strong relationship between the functional role of hubs and their participation in dynamic brain states; compared to other networks, the fronto-parietal network was found to demonstrate the greatest dynamic flexibility, as it is preferentially engaged in a wide variety of 64 different motor, cognitive, language, visual, and auditory tasks (Cole et al., 2013).

While the contributions of intrinsic functional connectivity networks have been widely established in their associations with cognition, research into how dynamic functional connectivity states contribute to cognition and intelligence is still developing. Recent work has demonstrated that aspects of these dynamic brain states are relevant to some aspects of cognition (Sadaghiani, Hesselmann, Friston, & Kleinschmidt, 2010; Thompson et al., 2013) in both healthy populations and a wide range of psychiatric and neurological disorders such as schizophrenia (Sakoglu et al., 2010), Alzheimer’s dementia (Jones et al., 2012), and Autism Spectrum Disorders (Starck et al., 2012). For example, some studies have shown that increases in DMN activity predicted error commission (Eichele et al., 2008), as well as temporary lapses in attention (Weissman, Roberts, Visscher, & Woldorff, 2006), which is supported by the countervailing role of the cingulo-opercular network in dynamic regulation of the DMN and sustained attention (Uddin, Kelly, Biswal, Castellanos, & Milham, 2009). Researchers have found that intra-individual differences in pre-stimulus network anti-correlation between the DMN and the fronto-parietal network was strongly predictive of both response time (Thompson et al., 2013) and response time variability (Kelly, Uddin, Biswal, Castellanos, & Milham, 2008). Other work has demonstrated that variability in the connectivity of the DMN is associated with more instances of off-task mind wandering (Kucyi & Davis, 2014). More recently, researchers found that while performing a cognitive control task, meta-stable states with strong links to cognitive control and visual networks increased in frequency relative to rest, while states linked to drowsiness and inattention decreased in frequency relative to rest (Hutchison & Morton, 2015). These findings suggest that dynamic states that emerge from intrinsic brain activity play a critical role in the attention and cognitive control processing that contributes to intelligence. We propose that the dynamic states that modulate these varied attention and cognitive control processes are directly linked to the cognitive procedures deployed while solving complex problems exemplified by either (i) inductive reasoning and fluid intelligence, or (ii) divergent thinking and creativity.

Overlapping Roles for Functional Brain Networks in Intelligence and Creativity

Traditional conceptualizations of the arts and sciences represented creativity and intelligence as separate domains, and the dominant trend in creativity research has supported this position, with meta-analyses showing very weak correlations between intelligence and creativity (Kim, 2005). More recently studies have demonstrated strong associations between creativity and intelligence (Plucker & Kaufman, 2011; Süß, Oberauer, Wittmann, Wilhelm, & Schulze, 2002). In most modern psychology research creativity is measured by the family of divergent thinking tasks (e.g., to generate uncommon uses of a “brick”), and although this is an important metric of creativity, most common psychometric methods of analyzing divergent thinking suffer from statistical constraints of unstable metrics of creativity and high collinearity of item uniqueness with item fluency (Nusbaum & Silvia, 2011). More recently, studies have investigated new ways of measuring creativity in divergent thinking tasks that do not suffer from these issues or sacrifice statistical power (Nusbaum & Silvia, 2011), and these analyses have revealed a much stronger and direct association between intelligence, executive function and creativity. For example, extensive evidence demonstrates that a range of executive functions are tied to creativity, such as working memory capacity (De Dreu, Nijstad, Baas, Wolsink, & Roskes, 2012; Süß et al., 2002), and both updating (Benedek, Jauk, Sommer, Arendasy, & Neubauer, 2014), and inhibition (Benedek, Franz, Heene, & Neubauer, 2012; Benedek et al., 2014). Furthermore, psychometric studies have demonstrated that general intelligence (Süß et al., 2002), and fluid intelligence are strongly coupled with performance in measurements of creativity such as divergent thinking originality (Süß et al., 2002), emotional metaphor creation (Silvia & Beaty, 2012), ideation originality (Benedek et al., 2012), and creative metaphor generation (Beaty & Silvia, 2013). These psychometric accounts are further buffeted by contemporary neuroimaging studies that detail further the mechanistic overlap between executive function and creativity.

Recent work has demonstrated that creativity and intelligence share an extensive array of overlapping neural correlates that center around the interaction between the default mode and fronto-parietal networks (Jung et al., 2013). The DMN plays an important role in internally generated thoughts that are both task-relevant and task-irrelevant, such as self-referential thought, autobiographical thoughts about the past and future (Andrews-Hanna, Smallwood, & Spreng, 2014; Christoff, Irving, Fox, Spreng, & Andrews-Hanna, 2016). Regions of the DMN are known to interact with hubs of the salience network, such as the anterior insula, and the fronto-parietal network, such as the dorsolateral prefrontal cortex (DLPFC). These executive networks are thought to play both a facilitating and constraining role on the DMN during creative tasks, for example, by engaging cognitive control processes that prevent interference from irrelevant self-generated items (Benedek et al., 2012). Beaty and colleagues found that high divergent thinking was explained by greater connectivity between the left inferior frontal gyrus and the entire DMN, and the right inferior frontal gyrus (IFG) showed greater connectivity to both the bilateral inferior parietal cortex and left DLPFC in the high creativity group as well (Beaty et al., 2014). In follow-up work, Beaty and colleagues found that global functional network efficiency was positively correlated with composite creativity scores, suggesting that greater efficiency of information transfer across networks is an important reflection of individual differences in creativity (Beaty et al., 2015). The importance of network connectivity in creativity has been underscored by studies demonstrating that white matter volume of the corpus callosum is positively correlated with creativity in the Torrance Tests of Creative Thinking, which may suggest that enhanced hemispheric specialization supports creative ideation through greater allowance for separate hemispheric processing (Moore et al., 2009). Furthermore, in a recent voxel-based lesion-symptom mapping study, researchers demonstrated that intelligence reliably predicted cognitive flexibility, that performance in these two factors both were dependent on a shared set of frontal, temporal, and parietal regions and white matter tracts (Barbey, Colom, & Grafman, 2013). More recently, intelligence researchers have begun experimentally probing the regions and networks involved in creativity. For example, one study found that in a divergent thinking task, cues to promote creative thought were associated with increased activity in the left frontopolar cortex and connectivity to the ACC (anterior cingulate cortex) and right frontopolar cortex (Green, Cohen, Raab, Yedibalian, & Gray, 2015). A creativity intervention study in which subjects were trained in divergent thinking found post-training behavioral improvements in response originality and fluency in untrained divergent thinking tasks (Sun et al., 2016). Furthermore, these changes were mirrored by group-level increases (post > pre) in functional activation in the bilateral dorsal ACC, DLPFC, and left inferior parietal lobe during an alternative uses task in the MRI scanner. This suggests that improving creative ability through training is both possible and dependent on regions of the brain critical to both cognitive control functions and self-generated thought.

Intelligence is thought to emerge through a dynamic hierarchical interaction between low level sensory regions, multimodal association regions in the parietal lobe, and frontal executive function regions such as the ACC and DLPFC (Jung & Haier 2007). We propose that creativity emerges through a similar pattern of dynamic hierarchical networks, with the default mode network playing a central role in generating a stream of internal stimuli that are fed forward to the parietal and frontal regions for abstraction, comparison, and response selection. Recent experimental work has provided the first evidence for such dynamic reconfiguration between the DMN, FPN and salience networks over the course of divergent thinking processing (Beaty et al., 2015). In this experiment, the authors split the time-course of response to a divergent thinking trial into five sections, and tested the dynamics of connectivity of seeds in the PCC, DLPFC, and precuneus. Their results suggest that at the beginning of the divergent thinking trial, the PCC first interacts with the bilateral insula, which is supported by prior work demonstrating the interaction between the insula and DMN (Uddin et al., 2009). The PCC continues by maintaining this connection and developing greater connectivity to frontal executive regions such as the DLPFC and ACC. Unlike the PCC, the right DLPFC does not demonstrate any initial changes in connectivity, but then increases in connectivity to the right PCC and inferior parietal lobe, which are both regions of the DMN. These results suggest that as is the case with executive function, in creativity the interaction between the salience network and the DMN is initially important, but in tests of creativity there are time-delayed increases in DMN-FPN coherence as the trial progresses, pointing to important similarities and contrasts between the dynamic network processing of intelligence and creativity.

Collectively these results demonstrate that intelligence and creativity share significant overlapping variance both psychometrically and mechanistically. Creativity, like many other higher cognitive attributes, is associated with intelligence at the level of both static and dynamic networks. In some sense, this is unsurprising given that human creativity and intelligence are regarded as evolutionarily highly advantageous abilities that would likely have emerged concurrently during human evolution. While these slow changes may have contributed to the emergence of intelligence and creativity over millennia, modern day children and adolescents provide us with other excellent models for the development of these higher cognitive functions. Understanding how these functional brain networks develop over the lifespan will give us important insights into the nature of creativity and intelligence.

## Neurodevelopmental Processes for Human Intelligence

Cognitive development during early childhood is dominated by critical periods of sensory and motor development while higher cognitive functions are largely undeveloped. The cognitive abilities most crucial to intelligence, such as cognitive control, fluid reasoning, and working memory, undergo substantial development during adolescence (Asato, Sweeney, & Luna, 2006; Huizinga, Dolan, & van der Molen, 2006). The onset of puberty leads to a cascade of hormonal changes that contribute to concurrent maturation of cognitive ability and brain structure (Crone & Dahl, 2012). This sudden onset manifests as an apparent imbalance in the development of regulatory competence to manage increases in arousal. In early and middle adolescence, pubertal onset enhances emotional arousal, reward sensitivity, and sensation seeking, and during middle adolescence low regulation of affect and cognition leads to vulnerability to risk taking and problem behavior (Steinberg, 2005). By late adolescence, regulatory competence is increased and risks are considerably lessened. Overall, adolescent cognitive development is marked by the creation of greater ability for self-directed and regulated cognition (i.e., cognitive control functions). Individuals show improvements in the capacity and efficiency of information processing as evidenced by increased reasoning ability. By late adolescence, individuals have already become more capable of complex, planned, abstract, hypothetical, and multidimensional thinking (Keating, Lerner, & Steinberg, 2004). The development of these cognitive abilities is central to the emergence of intelligence, and therefore the maturational trajectory of the associated brain regions, networks, and dynamic meta-stable states are critical markers not only of the development of adolescent cognition, but also of the emergence of intelligence specifically.

Brain development is strongly tied to a wide range of factors, including an organism’s physical environment (Greenough, Black, & Wallace, 1987), family, peers, nutrition, pubertal hormones, and education (Graber & Petersen, 1991). Intellectual stimulation during the first 12-30 months of life has a significant impact on a child’s future IQ (Carew, 1987), and types of motor and visual stimulation support development in object memory, discrimination, and recognition (Ruff, 1989; Schwarzer, 2014; Soska & Johnson, 2013; Spear, 2000). While network hubs are closely involved in cognitive performance, they likely play a critical role in facilitating development as well. Recent work has found that the rich club organization of the brain emerges very early during development, at only the 30^th^ week of gestation, followed by the integration of these rich club hubs with the rest of the brain (Ball et al., 2014). Furthermore, extensive changes in the size, myelination, and packing density of white matter axons during development contribute to an improvement in the efficacy of network communications (Paus, 2005; Paus et al., 1999). Thus the Network Dynamics Theory proposes that the development of white matter tracts assists in alleviating resource constraints on the communications infrastructure of child and adolescent brain networks, thereby enabling the concurrent emergence of adult phenotypic functional networks for intelligence.

While foundational intellectual development occurs during the first years of life, adolescence is the time period during which increases in most cognitive abilities occur (Graber & Petersen, 1991; Levin et al., 1991; Spear, 2000). Coincidentally, this is also the time period during which the PFC matures to its adult volume, with the sensorimotor, parietal and temporal cortices having already matured (Casey, Tottenham, Liston, & Durston, 2005). In adults the PFC plays a critical role in executive function and intelligence (Barbey et al., 2012; Barbey, Colom, Paul, & Grafman, 2013; Kane & Engle, 2002; Todd & Marois, 2005), and individual differences in the development of this brain region have been linked to differences in executive function in children, further supporting the role of the PFC in the development of intelligence (Casey et al., 1997). Prefrontal regions are underdeveloped during childhood, yet during this time they play an important role in cognitive abilities central to intelligence, such as cognitive control and working memory (Bunge, Dudukovic, Thomason, Vaidya, & Gabrieli, 2002; Durston, Thomas, Yang, Zimmerman, & Casey, 2002; Klingberg, Forssberg, & Westerberg, 2002). During childhood the prefrontal cortices increase in dendritic spine density, a critical marker of potential plasticity (Hering & Sheng, 2001), and this density peaks during human adolescence, reaching levels two or threefold greater than adult levels (Petanjek et al., 2011). The fronto-parietal network, which has been broadly implicated in cognitive control, working memory, and intelligence (Dosenbach et al., 2008; Jung & Haier, 2007; Nagy, Westerberg, & Klingberg, 2004), undergoes both gray matter and white matter maturation concurrently during development (Olesen, Nagy, Westerberg, & Klingberg, 2003). The maturation of this brain network parallels the development of intelligence as well, for during adolescence the long-range connections in this network strengthen and short-range connections weaken as it matures towards its adult form (Fair et al., 2007). Furthermore, recent evidence has demonstrated that adolescents also demonstrate greatest cortical development and white matter myelination in network hubs, which are critical for managing both internal and between network dynamics (Whitaker et al., 2016).

As static networks demonstrate long-term changes over time during adolescence associated with development of intelligence, long-term changes in dynamic brain networks serve as important markers of developmental trajectories as well (DiMartino et al., 2014; Hutchison & Morton, 2015); furthermore, the Network Dynamics Theory proposes that the dynamic connectome should be even more sensitive to developmental changes in intelligence than static networks. The neuroscience community has become increasingly interested in analyzing and characterizing brain dynamics in younger populations, and recent work by Hutchison and Morton (2015) has revealed several insightful differences between the dynamic connectomes of children and adults. In a study of 9 to 32 year olds, age was negatively associated with the number of brain states occupied during rest, meaning that adults switch between a more restricted set of brain states than children and teens. Furthermore, age was positively associated with the number of transitions between states and lower inter-transition intervals, meaning that adults not only switched between brain states more frequently, but when they began a transition they also completed it more rapidly. Given that the cognitive control networks are critical for switching between functional networks (Uddin, Supekar, Ryali, & Menon, 2011), and these cognitive control networks are less developed in children and adolescents (Power, Barnes, Snyder, Schlaggar, & Petersen, 2011; Supekar, Musen, & Menon, 2009), the Network Dynamics Theory proposes that these cognitive control meta-stable states are heavily involved in intelligence for their ability to regulate other brain networks, therefore allowing for more rapid information access, manipulation, and integration. This position is further supported by recent findings that adults demonstrate greater variability of network-to-network coupling compared to children and adolescents, and that this difference is most pronounced in cognitive control networks (Cole et al., 2013; Hutchison & Morton, 2015). We propose that adults are thus capable of using cognitive control networks in a greater variety of ways, and increasing efficiency of use may support the developmental emergence of higher cognitive functions that contribute to intelligence.

## Conclusions

Intelligence emerges through a set of extrinsically and intrinsically driven interactions. Dynamic brain networks interact with the extrinsic environment, which in turn drives the development of cognitive ability. Simultaneously, intrinsically driven developmental trajectories drive maturation of static brain networks and related dynamic network meta-stable states (Figure 1). We predict that while individual differences in static brain networks shed light on the development of intelligence from childhood to adulthood, concomitant changes in dynamic brain network meta-stable states should demonstrate even greater sensitivity to the development of cognitive abilities associated with both intelligence and creativity. Specifically, Network Dynamics Theory predicts that increasing network efficiency and modularity in dynamic states of the rich club and fronto-parietal networks over development should mirror the improvements in executive and other higher cognitive functions associated with intelligence. Furthermore, we predict that as children and teens mature, the increasing speed of switching between dynamic meta-stable brain states should be a critical marker for the development of intelligence and creativity (DiMartino et al., 2014; Hutchison & Morton, 2015).

Network Dynamics Theory has important implications for both the study of intelligence and the efforts underway to improve intelligence and creativity (Table 2). Given that intelligence improves via development of dynamic network states during adolescence, we predict that interventions aimed at training executive functions during adolescence may see particular success in changing the dynamic brain states that accompany the development of intelligence. This is particularly supported by findings that the PFC is highly plastic during adolescence (Petanjek et al., 2011), suggesting that during adolescence executive processing may be subject to greater experience-induced plasticity. More specifically, we hypothesize that training on executive function tasks during adolescence may change the processing style of these tasks, training adolescents to engage in faster switching between brain states, which may be employed in other scenarios as well. Generalization of training may thus be more accurately detected through changes in the dynamic meta-stable states from which intelligence and creativity emerge.

**Table 2.** This table summarizes the key predictions made by the Network Dynamics Theory of intelligence regarding the role of specific brain networks and development in intelligence. ACC = anterior cingulate cortex; DLPFC = dorsolateral prefrontal cortex.

|  |
| --- |
| 1. Adults have a greater capacity to use cognitive control networks to support a variety of cognitive tasks (relative to adolescents), and this increasing efficiency of use supports the developmental emergence of higher cognitive functions that contribute to intelligence |
| 2. The dynamic connectome provides a more sensitive and specific marker of developmental changes in intelligence than classic approaches that focus on a single network or static network properties |
| 3. Both intelligence and creativity depend on the emergence of macro-level network patterns, such as global efficiency and rapid meta-stable state switching, in response to both external and internal stimuli |
| 4. The maturation of macro-level topological properties of the rich club network plays a central role in human intelligence |
| 5. Dynamic brain states that modulate cognitive control support intelligence and creativity through their guidance of the cognitive procedures deployed while solving complex problems |
| 6. The increase in fronto-parietal network efficiency and modularity over development are predicted to mirror the improvements in executive and other higher cognitive functions associated with intelligence |
| 7. Interventions aimed at training executive functions during adolescence are predicted to change the dynamic interactions between the fronto-parietal and default mode networks that accompany the development of intelligence and creativity |
| 8. Training on executive function tasks during adolescence will enable domain-general learning strategies that may be applied in new scenarios. For example, training that encourages adolescents to engage in faster switching between brain states may be more likely to demonstrate transfer across a broad spectrum of cognitive control tasks |
| 9. Creativity emerges through a dynamic hierarchical interaction between default mode regions, multimodal association regions in the parietal lobe, and frontal executive regions such as the ACC and DLPFC |

## References

Allen, E. A., Damaraju, E., Plis, S. M., Erhardt, E. B., Eichele, T., & Calhoun, V. D. (2014). Tracking whole-brain connectivity dynamics in the resting state. Cerebral Cortex, 24(3), 663–676. http://doi.org/10.1093/cercor/bhs352

Andrews-Hanna, J. R., Smallwood, J., & Spreng, R. N. (2014). The default network and self-generated thought: component processes, dynamic control, and clinical relevance. Annals of the New York Academy of Sciences, 1316(1), 29–52. http://doi.org/10.1111/nyas.12360

Asato, M. R., Sweeney, J. A., & Luna, B. (2006). Cognitive processes in the development of TOL performance. Neuropsychologia, 44(12), 2259–2269. http://doi.org/10.1016/j.neuropsychologia.2006.05.010

Ball, G., Aljabar, P., Zebari, S., Tusor, N., Arichi, T., Merchant, N., … Counsell, S. J. (2014). Rich-club organization of the newborn human brain. Proceedings of the National Academy of Sciences of the United States of America, 111(20), 7456–61. http://doi.org/10.1073/pnas.1324118111

Barbey, A. K., Colom, R., & Grafman, J. (2013a). Architecture of cognitive flexibility revealed by lesion mapping. NeuroImage, 82, 547–554. http://doi.org/10.1016/j.neuroimage.2013.05.087

Barbey, A. K., Colom, R., & Grafman, J. (2013b). Dorsolateral prefrontal contributions to human intelligence. Neuropsychologia, 51, 1361–1369. http://dx.doi.org/10.1016/j.neuropsychologia.2012.05.017

Barbey, A. K., Colom, R., Solomon, J., Krueger, F., Forbes, C., & Grafman, J. (2012). An integrative architecture for general intelligence and executive function revealed by lesion mapping. Brain: A Journal of Neurology, 135(Pt 4), 1154–64. http://doi.org/10.1093/brain/aws021

Beaty, R. E., Benedek, M., Barry Kaufman, S., & Silvia, P. J. (2015). Default and Executive Network Coupling Supports Creative Idea Production. Scientific Reports, 5, 10964. http://doi.org/10.1038/srep10964

Beaty, R. E., Benedek, M., Wilkins, R. W., Jauk, E., Fink, A., Silvia, P. J., … Neubauer, A. C. (2014). Creativity and the default network: A functional connectivity analysis of the creative brain at rest. Neuropsychologia, 64C, 92–98. http://doi.org/10.1016/j.neuropsychologia.2014.09.019

Beaty, R. E., & Silvia, P. J. (2013). Metaphorically speaking: cognitive abilities and the production of figurative language, 255–267. http://doi.org/10.3758/s13421-012-0258-5

Benedek, M., Franz, F., Heene, M., & Neubauer, A. C. (2012). Differential effects of cognitive inhibition and intelligence on creativity. Personality and Individual Differences, 53(4), 480–485. http://doi.org/10.1016/j.paid.2012.04.014

Benedek, M., Jauk, E., Sommer, M., Arendasy, M., & Neubauer, A. C. (2014). Intelligence, creativity, and cognitive control: The common and differential involvement of executive functions in intelligence and creativity. Intelligence, 46(1), 73–83. http://doi.org/10.1016/j.intell.2014.05.007

Braver, T., Cohen, J., Nystrom, L., & Jonides, J. (1997). A parametric study of prefrontal cortex involvement in human working memory. Neuroimage, (5), 49–62. Retrieved from http://www.sciencedirect.com/science/article/pii/S1053811996902475

Bullmore, E., & Sporns, O. (2009). Complex brain networks: graph theoretical analysis of structural and functional systems. Nat Rev Neurosci, 10(3), 186–198. Journal Article. http://doi.org/10.1038/nrn2575

Bullmore, E. T., & Bassett, D. S. (2011). Brain Graphs: Graphical Models of the Human Brain Connectome. Annu. Rev. Clin. Psychol, 7(1), 113–40. Journal Article. http://doi.org/10.1146/annurev-clinpsy-040510-143934

Bunge, S. A., Dudukovic, N. M., Thomason, M. E., Vaidya, C. J., & Gabrieli, J. D. E. (2002). Immature Frontal Lobe Contributions to Cognitive Control in Children: Evidence from fMRI, 33, 301–311.

Byrge, L., Sporns, O., & Smith, L. B. (2014). Developmental process emerges from extended brain – body – behavior networks. Trends in Cognitive Sciences, 18(8), 395–403. http://doi.org/10.1016/j.tics.2014.04.010

Callicott, J. H., Mattay, V. S., Bertolino, A., Finn, K., Coppola, R., Frank, J. A., … Weinberger, D. R. (1999). Physiological characteristics of capacity constraints in working memory as revealed by functional MRI. Cereb Cortex, 9(1), 20–26. Retrieved from http://www.ncbi.nlm.nih.gov/pubmed/10022492

Carew, J. V. (1987). Experience and the development of intelligence in young children at home and in day care. Monographs of the Society for Research in Child Development (Vol. 45).

Casey, B. J., Castellanos, F. X., Giedd, J. N., Marsh, W. L., Hamburger, S. D., Schubert, A. B., … Rapoport, J. L. (1997). Implication of Right Frontostriatal Circuitry in Response Inhibition and Attention-Deficit/Hyperactivity Disorder. Journal of the American Academy of Child & Adolescent Psychiatry, 36(3), 374–383. http://doi.org/10.1097/00004583-199703000-00016

Casey, B. J., Tottenham, N., Liston, C., & Durston, S. (2005). Imaging the developing brain: what have we learned about cognitive development?, 9(3). http://doi.org/10.1016/j.tics.2005.01.011

Christoff, K., Irving, Z. C., Fox, K. C. R., Spreng, R. N., & Andrews-Hanna, J. R. (2016). Mind-wandering as spontaneous thought: a dynamic framework. Nature Reviews Neuroscience, 1–44. http://doi.org/10.1038/nrn.2016.113

Cole, M. W., Reynolds, J. R., Power, J. D., Repovs, G., Anticevic, A., & Braver, T. S. (2013). Multi-task connectivity reveals flexible hubs for adaptive task control. Nature Neuroscience, 16(9), 1348–1355. http://doi.org/10.1038/nn.3470

Crone, E. A., & Dahl, R. E. (2012). Understanding adolescence as a period of social – affective engagement and goal flexibility. Nature, 13(9), 636–650. http://doi.org/10.1038/nrn3313

Crossley, N. A., Mechelli, A., Scott, J., Carletti, F., Fox, P. T., Mcguire, P., & Bullmore, E. T. (2014). The hubs of the human connectome are generally implicated in the anatomy of brain disorders. Brain, 137(8), 2382–2395. http://doi.org/10.1093/brain/awu132

Crossley, N. A., Mechelli, A., Vértes, P. E., Winton-Brown, T. T., Patel, A. X., Ginestet, C. E., … Bullmore, E. T. (2013). Cognitive relevance of the community structure of the human brain functional coactivation network. Proceedings of the National Academy of Sciences of the United States of America, 110(28), 11583–11588. http://doi.org/10.1073/pnas.1220826110

De Dreu, C. K. W., Nijstad, B. A., Baas, M., Wolsink, I., & Roskes, M. (2012). Working memory benefits creative insight, musical improvisation, and original ideation through maintained task-focused attention. Personality and Social Psychology Bulletin, 38(5), 656–669. http://doi.org/10.1177/0146167211435795

Deco, G., Jirsa, V. K., & McIntosh, A. R. (2011). Emerging concepts for the dynamical organization of resting-state activity in the brain. Nature Reviews. Neuroscience, 12(1), 43–56. http://doi.org/10.1038/nrn2961

Dehaene, S., & Changeux, J. P. (2011). Experimental and Theoretical Approaches to Conscious Processing. Neuron, 70(2), 200–227. http://doi.org/10.1016/j.neuron.2011.03.018

Dehaene, S., Kerszberg, M., & Changeux, J. P. (1998). A neuronal model of a global workspace in effortful cognitive tasks. Proceedings of the National Academy of Sciences of the United States of America, 95(24), 14529–14534. http://doi.org/10.1073/pnas.95.24.14529

DiMartino, A., Fair, D. A., Kelly, C., Satterthwaite, T. D., Castellanos, F. X., Thomason, M. E., … Milham, M. P. (2014). Unraveling the miswired connectome: A developmental perspective. Neuron, 83(6), 1335–1353. http://doi.org/10.1016/j.neuron.2014.08.050

Dosenbach, N. U. F., Fair, D. A., Cohen, A. L., Schlaggar, B. L., & Petersen, S. E. (2008). A dual-networks architecture of top-down control. Trends in Cognitive Sciences, 12(3), 99–105. http://doi.org/10.1016/j.tics.2008.01.001

Dosenbach, N. U. F., Visscher, K. M., Palmer, E. D., Miezin, F. M., Wenger, K. K., Kang, H. C., … Petersen, S. E. (2006). A core system for the implementation of task sets. Neuron, 50(5), 799–812. http://doi.org/10.1016/j.neuron.2006.04.031

Duncan, J. (2010). The multiple-demand (MD) system of the primate brain: mental programs for intelligent behaviour. Trends in Cognitive Sciences, 14(4), 172–9. http://doi.org/10.1016/j.tics.2010.01.004

Durston, S., Thomas, K. M., Yang, Y., Zimmerman, R. D., & Casey, B. J. (2002). A neural basis for the development of inhibitory control. Developmental Science, 4, 9–16.

Eichele, T., Debener, S., Calhoun, V. D., Specht, K., Engel, A. K., Hugdahl, K., … Ullsperger, M. (2008). Prediction of human errors by maladaptive changes in eventrelated brain networks. Proceedings of the National Academy of Sciences of the United States of America, 105(16), 6173–8. http://doi.org/10.1073/pnas.0708965105

Fair, D. A., Dosenbach, N. U. F., Church, J. A., Cohen, A. L., Brahmbhatt, S., Miezin, F. M., … Schlaggar, B. L. (2007). Development of distinct control networks through segregation and integration. Proceedings of the National Academy of Sciences of the United States of America, 104, 13507–13512. http://doi.org/10.1073/pnas.0705843104

Finn, E. S., Shen, X., Scheinost, D., Rosenberg, M. D., Huang, J., Chun, M. M., … Constable, R. T. (2015). Functional connectome fingerprinting: identifying individuals using patterns of brain connectivity. Nature Neuroscience, (October), 1–11. http://doi.org/10.1038/nn.4135

Gazzaniga, M. S. (2000). Cerebral specialization and interhemispheric communication: does the corpus callosum enable the human condition? Brain: A Journal of Neurology, 123 (Pt 7, 1293–1326. http://doi.org/10.1093/brain/123.7.1293

Giessing, C., Thiel, C. M., Alexander-Bloch, A. F., Patel, A. X., & Bullmore, E. T. (2013). Human Brain Functional Network Changes Associated with Enhanced and Impaired Attentional Task Performance. Journal of Neuroscience, 33(14), 5903–5914. http://doi.org/10.1523/JNEUROSCI.4854-12.2013

Gläscher, J., Rudrauf, D., Colom, R., Paul, L. K., Tranel, D., Damasio, H., & Adolphs, R. (2010). Distributed neural system for general intelligence revealed by lesion mapping\r10.1073/pnas.0910397107. Proceedings of the National Academy of Sciences, 107(10), 4705–4709. http://doi.org/10.1073/pnas.0910397107

Graber, J. A., & Petersen, A. C. (1991). Cognitive changes at adolescence: Biological perspectives. Foundations of human behavior [Brain maturation and cognitive development: Comparative and cross-cultural perspectives]. Retrieved from http://www.sciencedirect.com/science/article/B6WVC-446CXBY-1WY/1/6dab32f6d1b86b9e94ff7d3c6262614c

Green, A. E., Cohen, M. S., Raab, H. A., Yedibalian, C. G., & Gray, J. R. (2015). Frontopolar activity and connectivity support dynamic conscious augmentation of creative state. Human Brain Mapping, 36(3), 923–934. http://doi.org/10.1002/hbm.22676

Greenough, W. T., Black, J. E., & Wallace, C. S. (1987). Experience and brain development. Child Development, 58(3), 539–559. http://doi.org/10.2307/1130197

Güntürkün, O. (2005). Avian and mammalian “prefrontal cortices”: limited degrees of freedom in the evolution of the neural mechanisms of goal-state maintenance. Brain Research Bulletin, 66(4–6), 311–6. http://doi.org/10.1016/j.brainresbull.2005.02.004

Hampshire, A., Highfield, R. R., Parkin, B. L., & Owen, A. M. (2012). Fractionating Human Intelligence. Neuron, 76(6), 1225–1237. http://doi.org/10.1016/j.neuron.2012.06.022

Harriger, L., van den Heuvel, M. P., & Sporns, O. (2012). Rich Club Organization of Macaque Cerebral Cortex and Its Role in Network Communication. PLoS ONE, 7(9). http://doi.org/10.1371/journal.pone.0046497

Hering, H., & Sheng, M. (2001). Dendritic spines: structure, dynamics and regulation. Nature Reviews. Neuroscience, 2(12), 880–888. http://doi.org/10.1038/35104061

Huizinga, M., Dolan, C. V., & van der Molen, M. W. (2006). Age-related change in executive function: Developmental trends and a latent variable analysis. Neuropsychologia, 44(11), 2017–2036. http://doi.org/10.1016/j.neuropsychologia.2006.01.010

Hutchison, R. M., & Morton, J. B. (2015). Tracking the Brain’s Functional Coupling Dynamics over Development. The Journal of Neuroscience, 35(17), 6849–6859. http://doi.org/10.1523/JNEUROSCI.4638-14.2015

Hutchison, R. M., Womelsdorf, T., Allen, E. A., Bandettini, P. A., Calhoun, V. D., Corbetta, M., … Chang, C. (2013). Dynamic functional connectivity: Promise, issues, and interpretations. NeuroImage, 80, 360–378. http://doi.org/10.1016/j.neuroimage.2013.05.079

Jaeggi, S. M., Buschkuehl, M., Etienne, A., Ozdoba, C., Perrig, W. J., & Nirkko, A. C. (2007). On how high performers keep cool brains in situations of cognitive overload. Cognitive, Affective & Behavioral Neuroscience, 7(2), 75–89. http://doi.org/10.3758/CABN.7.2.75

Jones, D. T., Vemuri, P., Murphy, M. C., Gunter, J. L., Senjem, M. L., Machulda, M. M., … Jack, C. R. (2012). Non-stationarity in the “resting brain’s” modular architecture. PLoS ONE, 7(6). http://doi.org/10.1371/journal.pone.0039731

Jung, R. E., Brooks, W. M., Yeo, R. A., Chiulli, S. J., Weers, D. C., & Sibbitt, W. L. (1999). Biochemical markers of intelligence: a proton MR spectroscopy study of normal human brain. Proceedings. Biological Sciences / The Royal Society, 266(1426), 1375–9. http://doi.org/10.1098/rspb.1999.0790

Jung, R. E., & Haier, R. J. (2007). The Parieto-Frontal Integration Theory (P-FIT) of intelligence: converging neuroimaging evidence. The Behavioral and Brain Sciences, 30(2), 135–54-87. http://doi.org/10.1017/S0140525X07001185

Jung, R. E., Haier, R. J., Yeo, R. A., Rowland, L. M., Petropoulos, H., Levine, A. S., … Brooks, W. M. (2005). Sex differences in N-acetylaspartate correlates of general intelligence: An 1H-MRS study of normal human brain. NeuroImage, 26(3), 965–972. http://doi.org/10.1016/j.neuroimage.2005.02.039

Just, M. A., Carpenter, P. A., Maguire, M., Diwadkar, V., & McMains, S. (2001). Mental rotation of objects retrieved from memory: a functional MRI study of spatial processing. Journal of Experimental Psychology. General, 130(3), 493–504. http://doi.org/10.1037/0096-3445.130.3.493

Just, M. A., & Varma, S. (2007). The organization of thinking: what functional brain imaging reveals about the neuroarchitecture of complex cognition. Cognitive, Affective & Behavioral Neuroscience, 7(3), 153–191. http://doi.org/10.3758/CABN.7.3.153

Kane, M. J., & Engle, R. W. (2002). The role of prefrontal cortex in working-memory capacity, executive attention, and general fluid intelligence: an individual-differences perspective. Psychonomic Bulletin & Review, 9(4), 637–71. Retrieved from http://www.ncbi.nlm.nih.gov/pubmed/12613671

Keating, D. P., Lerner, R. M., & Steinberg, L. (2004). Cognitive and brain development. Handbook of adolescent psychology (Vol. 2). http://doi.org/10.4074/S0013754512003035

Kelly, A. M. C., Uddin, L. Q., Biswal, B. B., Castellanos, F. X., & Milham, M. P. (2008). Competition between functional brain networks mediates behavioral variability. NeuroImage, 39(1), 527–537. http://doi.org/10.1016/j.neuroimage.2007.08.008

Kim, K. H. (2005). Can only intelligent people be creative? A meta-analysis. Journal of Secondary Gifted Education, 16(2–3), 57–66. http://doi.org/10.4219/jsge-2005-473

Kitzbichler, M. G., Henson, R. N. A., Smith, M. L., Nathan, P. J., & Bullmore, E. T. (2011). Cognitive Effort Drives Workspace Configuration of Human Brain Functional Networks. Journal of Neuroscience, 31(22), 8259–8270. http://doi.org/10.1523/JNEUROSCI.0440-11.2011

Klingberg, T., Forssberg, H., & Westerberg, H. (2002). Training of working memory in children with ADHD. J Clin Exp Neuropsychol, 24(6), 781–791. http://doi.org/10.1076/jcen.24.6.781.8395

Kucyi, A., & Davis, K. D. (2014). Dynamic functional connectivity of the default mode network tracks daydreaming. NeuroImage, 100, 471–480. http://doi.org/10.1016/j.neuroimage.2014.06.044

Langeslag, S. J. E., Schmidt, M., Ghassabian, A., Jaddoe, V. W., Hofman, A., Lugt, A. Van Der, … White, T. J. H. (2012). Functional Connectivity between Parietal and Frontal Brain Regions and Intelligence in Young Children: The Generation R Study, *0*(May). http://doi.org/10.1002/hbm.22143

Levin, H. S., Culhane, K. a., Hartmann, J., Evankovich, K., Mattson, A. J., Harward, H., … Fletcher, J. M. (1991). Developmental changes in performance on tests of purported frontal lobe functioning. Developmental Neuropsychology, 7(June 2014), 377–395. http://doi.org/10.1080/87565649109540499

Liu, X., Chang, C., & Duyn, J. H. (2013). Decomposition of spontaneous brain activity into distinct fMRI co-activation patterns. Frontiers in Systems Neuroscience, 7(December), 101. http://doi.org/10.3389/fnsys.2013.00101

Miller, E. K., & Cohen, J. D. (2001). An integrative theory of prefrontal cortex function. Annual Review of Neuroscience, 24(1), 167–202. Retrieved from http://www.ncbi.nlm.nih.gov/pubmed/11283309

Moore, D. W., Bhadelia, R. A., Billings, R. L., Fulwiler, C., Heilman, K. M., Rood, K. M. J., & Gansler, D. A. (2009). Hemispheric connectivity and the visual-spatial divergent-thinking component of creativity. Brain and Cognition, 70(3), 267–272. http://doi.org/10.1016/j.bandc.2009.02.011

Moussa, M. N. M., Vechlekar, C. D. C., Burdette, J. H., Steen, M. R., Hugenschmidt, C. E., & Laurienti, P. J. (2011). Changes in Cognitive State Alter Human Functional Brain Networks. Frontiers in Human Neuroscience, 5(August), 1–15. http://doi.org/10.3389/fnhum.2011.00083

Nagy, Z., Westerberg, H., & Klingberg, T. (2004). Maturation of white matter is associated with the development of cognitive functions during childhood. J Cogn Neurosci, 16(7), 1227–1233. http://doi.org/10.1162/0898929041920441 [doi]http://doi.org/10.1162/0898929041920441 [doi]

Newman, S. D., & Just, M. A. (2005). The Neural Bases of Intelligence: A Perspective Based on Functional Neuroimaging. Cognition and Intelligence, 88–103. http://doi.org/10.1017/CBO9780511607073.006

Nikolaidis, A., Baniqued, P. L., Kranz, M. B., Scavuzzo, C. J., Barbey, A. K., Kramer, A. F., & Larsen, R. J. (2016). Multivariate Associations of Fluid Intelligence and NAA. Cerebral Cortex, bhw070. http://doi.org/10.1093/cercor/bhw070

Nikolaidis, A., Goatz, D., Smaragdis, P., & Kramer, A. (2015). Predicting Skill-Based Task Performance and Learning with fMRI Motor and Subcortical Network Connectivity. 2015 International Workshop on Pattern Recognition in NeuroImaging, 93–96. IEEE. http://doi.org/10.1109/PRNI.2015.35

Nikolaidis, A., Voss, M. W., Lee, H., Vo, L. T. K., & Kramer, A. F. (2014). Parietal plasticity after training with a complex video game is associated with individual differences in improvements in an untrained working memory task. Frontiers in Human Neuroscience, 8(March), 1–11. http://doi.org/10.3389/fnhum.2014.00169

Nusbaum, E. C., & Silvia, P. J. (2011). Are intelligence and creativity really so different?. Fluid intelligence, executive processes, and strategy use in divergent thinking. Intelligence, 39(1), 36–45. http://doi.org/10.1016/j.intell.2010.11.002

Olesen, P. J., Nagy, Z., Westerberg, H., & Klingberg, T. (2003). Combined analysis of DTI and fMRI data reveals a joint maturation of white and grey matter in a frontoparietal network. Cognitive Brain Research, 18(1), 48–57. http://doi.org/10.1016/j.cogbrainres.2003.09.003

Pascual-Leone, A., Amedi, A., Fregni, F., & Merabet, L. B. (2005). The Plastic Human Brain Cortex. Annual Review of Neuroscience, 28(1), 377–401. http://doi.org/10.1146/annurev.neuro.27.070203.144216

Paul, E. J., Larsen, R. J., Nikolaidis, A., Ward, N., Hillman, C. H., Cohen, N. J., … Barbey, A. K. (2016). Dissociable Brain Biomarkers of Fluid Intelligence. NeuroImage, (May). http://doi.org/10.1016/j.neuroimage.2016.05.037

Paus, T. (2005). Mapping brain maturation and cognitive development during adolescence. Trends in Cognitive Sciences; Trends in Cognitive Sciences.

Paus, T., Zijdenbos, A., Worsley, K., Collins, D. L., Blumenthal, J., Giedd, J. N., … Evans, A. C. (1999). Structural maturation of neural pathways in children and adolescents: in vivo study. Science (New York, N.Y.), 283(5409), 1908–1911. http://doi.org/10.1126/science.283.5409.1908

Penke, L., Maniega, S. M., Bastin, M. E., Valdés Hernández, M. C., Murray, C., Royle, N. A., … Deary, I. J. (2012). Brain white matter tract integrity as a neural foundation for general intelligence. Molecular Psychiatry, 17(10), 1026–1030. http://doi.org/10.1038/mp.2012.66"

Petanjek, Z., Judas, M., Simic, G., Rasin, M. R., Uylings, H. B. M., Rakic, P., & Kostovic, I. (2011). Extraordinary neoteny of synaptic spines in the human prefrontal cortex. Proceedings of the National Academy of Sciences of the United States of America, 108(32), 13281–13286. http://doi.org/10.1073/pnas.1105108108

Plucker, J. A., & Kaufman, J. C. (2011). Intelligence and Creativity. The Cambridge Handbook of Intelligence, (4), 771–783. http://doi.org/10.1037/e518652004-001

Power, J. D., Barnes, K. A., Snyder, A. Z., Schlaggar, B. L., & Petersen, S. E. (2011). NeuroImage Spurious but systematic correlations in functional connectivity MRI networks arise from subject motion. NeuroImage. http://doi.org/10.1016/j.neuroimage.2011.10.018

Power, J. D., Cohen, A. L., Nelson, S. M., Wig, G. S., Barnes, K. A., Church, J. a, … Petersen, S. E. (2011). Functional network organization of the human brain. Neuron, 72(4), 665–78. http://doi.org/10.1016/j.neuron.2011.09.006

Power, J. D., & Petersen, S. E. (2013). Control-related systems in the human brain. Current Opinion in Neurobiology, 23(2), 223–8. http://doi.org/10.1016/j.conb.2012.12.009

Röder, B., Stock, O., Neville, H., Bien, S., & Rösler, F. (2002). Brain activation modulated by the comprehension of normal and pseudo-word sentences of different processing demands: a functional magnetic resonance imaging study. NeuroImage, 15(4), 1003–1014. http://doi.org/10.1006/nimg.2001.1026

Ross, A. J., & Sachdev, P. S. (2004). Magnetic resonance spectroscopy in cognitive research. Brain Research. Brain Research Reviews, 44(2–3), 83–102. http://doi.org/10.1016/j.brainresrev.2003.11.001

Roth, G., & Dicke, U. (2005). Evolution of the brain and intelligence. Trends in Cognitive Sciences, 9(5), 250–257. http://doi.org/10.1016/j.tics.2005.03.005

Rubinov, M., & Sporns, O. (2009). NeuroImage Complex network measures of brain connectivity: Uses and interpretations. NeuroImage, 52(3), 1059–1069. http://doi.org/10.1016/j.neuroimage.2009.10.003

Ruff, H. A. (1989). The infant’s use of visual and haptic information in the perception and recognition of objects. Canadian Journal of Psychology, 43, 302–319. http://doi.org/10.1037/h0084222

Sabaté, M., González, B., & Rodríguez, M. (2004). Brain lateralization of motor imagery: Motor planning asymmetry as a cause of movement lateralization. Neuropsychologia, 42(8), 1041–1049. http://doi.org/10.1016/j.neuropsychologia.2003.12.015

Sadaghiani, S., Hesselmann, G., Friston, K. J., & Kleinschmidt, A. (2010). The relation of ongoing brain activity, evoked neural responses, and cognition. Frontiers in Systems Neuroscience, 4(June), 20. http://doi.org/10.3389/fnsys.2010.00020

Sakoglu, U., Pearlson, G. D., Kiehl, K. A., Wang, Y. M., Michael, A. M., & Calhoun, V. D. (2010). A method for evaluating dynamic functional network connectivity and task-modulation: Application to schizophrenia. Magnetic Resonance Materials in Physics, Biology and Medicine, 23(5–6), 351–366. http://doi.org/10.1007/s10334-010-0197-8

Schwarzer, G. (2014). How Motor and Visual Experiences Shape Infants’ Visual Processing of Objects and Faces. Child Development Perspectives, 8(4), 213–217. http://doi.org/10.1111/cdep.12093

Silvia, P. J., & Beaty, R. E. (2012). Making creative metaphors: The importance of fluid intelligence for creative thought. Intelligence, 40(4), 343–351. http://doi.org/10.1016/j.intell.2012.02.005

Soska, K. C., & Johnson, S. P. (2013). Development of Three-Dimensional Completion of Complex Objects. Infancy, 18(3), 325–344. http://doi.org/10.1111/j.1532-7078.2012.00127.x

Spear, L. P. (2000). The adolescent brain and age-related behavioral manifestations. Neuroscience and Biobehavioral Reviews, 24(4), 417–463. http://doi.org/10.1016/S0149-7634(00)00014-2

Starck, T., Nikkinen, J., Remes, J., Rahko, J., Moilanen, I., Tervonen, O., & Kiviniemi, V. (2012). Temporally varying connectivity between ICA default-mode sub-networks—ASD vs. controls. In Organization for Human Brain Mapping. Beijing.

Steinberg, L. (2005). Cognitive and affective development in adolescence. Trends in Cognitive Sciences. http://doi.org/10.1016/j.tics.2004.12.005

Sun, J., Chen, Q., Zhang, Q., Li, Y., Li, H., Wei, D., … Qiu, J. (2016). Training your brain to be more creative: Brain functional and structural changes induced by divergent thinking training. Human Brain Mapping, (May). http://doi.org/10.1002/hbm.23246

Supekar, K., Musen, M., & Menon, V. (2009). Development of Large-Scale Functional Brain Networks in Children, 7(7). http://doi.org/10.1371/journal.pbio.1000157

Süß, H. M., Oberauer, K., Wittmann, W. W., Wilhelm, O., & Schulze, R. (2002). Working-memory capacity explains reasoning ability - And a little bit more. Intelligence, 30(3), 261–288. http://doi.org/10.1016/S0160-2896(01)00100-3

Thompson, G. J., Magnuson, M. E., Merritt, M. D., Schwarb, H., Pan, W. J., Mckinley, A., … Keilholz, S. D. (2013). Short-time windows of correlation between large-scale functional brain networks predict vigilance intraindividually and interindividually. Human Brain Mapping, 34(12), 3280–3298. http://doi.org/10.1002/hbm.22140

Todd, J. J., & Marois, R. (2005). Posterior parietal cortex activity predicts individual differences in visual short-term memory capacity. Cognitive, Affective, & Behavioral Neuroscience, 5(2), 144–155. http://doi.org/10.3758/CABN.5.2.144

Uddin, L. Q., Kelly, A. M. C., Biswal, B. B., Castellanos, F. X., & Milham, M. P. (2009). Functional Connectivity of Default Mode Network Components: Correlation, Anticorrelation, and Causality. Human Brain Mapping, 30(2), 625–637. http://doi.org/10.1002/hbm.20531

Uddin, L. Q., Supekar, K. S., Ryali, S., & Menon, V. (2011). Dynamic reconfiguration of structural and functional connectivity across core neurocognitive brain networks with development. J Neurosci, 31(50), 18578–18589. http://doi.org/10.1523/JNEUROSCI.4465-11.2011

Vakhtin, A. A., Ryman, S. G., Flores, R. A., & Jung, R. E. (2014). Functional brain networks contributing to the Parieto-Frontal Integration Theory of Intelligence. NeuroImage, 103, 349–354. http://doi.org/10.1016/j.neuroimage.2014.09.055

van den Heuvel, M. P., Kahn, R. S., Goñi, J., & Sporns, O. (2012). High-cost, highcapacity backbone for global brain communication. Proceedings of the National Academy of Sciences of the United States of America, 109(28), 11372–77. http://doi.org/10.1073/pnas.1203593109/-/DCSupplemental.www.pnas.org/cgi/doi/10.1073/pnas.1203593109

van den Heuvel, M. P., & Sporns, O. (2013). Network hubs in the human brain. Trends in Cognitive Sciences, 17(12), 683–696. http://doi.org/10.1016/j.tics.2013.09.012

van den Heuvel, M. P., Stam, C. J., Kahn, R. S., & Hulshoff Pol, H. E. (2009). Efficiency of functional brain networks and intellectual performance. The Journal of Neuroscience: The Official Journal of the Society for Neuroscience, 29(23), 7619–7624. http://doi.org/10.1523/JNEUROSCI.1443-09.2009

Weissman, D. H., Roberts, K. C., Visscher, K. M., & Woldorff, M. G. (2006). The neural bases of momentary lapses in attention. Nature Neuroscience, 9(7), 971–8. http://doi.org/10.1038/nn1727

Whitaker, K. J., Vértes, P. E., Romero-Garcia, R., Váša, F., Moutoussis, M., Prabhu, G., … Bullmore, E. T. (2016). Adolescence is associated with genomically patterned consolidation of the hubs of the human brain connectome. Proceedings of the National Academy of Sciences, 201601745. http://doi.org/10.1073/pnas.1601745113

